# Tunneling Effect with Time-Dependent Effective Potential Barrier: A Semiclassical (WKB) Reinterpretation of Drug Release Kinetics in Polymeric Nanocapsules

**DOI:** 10.64898/2026.07.30.741891

**Authors:** Douglas F. de Albuquerque, Maria Antonia S. de Albuquerque

**Author notes:** Corresponding author (D.F.d.A.); (M.A.S.d.A.).

## Abstract

Recent models describe drug release from polymeric nanoparticles through an analogy with the quantum tunneling effect, treating the delivery system as a static rectangular potential barrier. In this work, we argue that this analogy is structurally identical to the standard solution of the Schrödinger equation for a rectangular barrier, and that the introduction of a multifractal formalism to describe time evolution — obtained via a formal Wick rotation (*x* → *t*) — lacks direct physical justification. We propose, instead, to treat the barrier height as an effective function of time, *U*_eff_(*t*) = *U*_0_ *f*(*t*), with *f*(*t*) varying slowly within the barrier region, reflecting the progressive degradation/swelling of the polymeric matrix, under two hypotheses for *f*(*t*)— exponential decay and rational decay (Hill-type). Rather than the thick-barrier WKB approximation, the *exact* transmission formula is used throughout, which is real-analytic in *f*(*t*) and continues smoothly into the resonance (over-barrier) regime once the barrier collapses, avoiding the artificial step-like transitions produced by the WKB approximation used in earlier drafts of this work. Both hypotheses for *f*(*t*) predict a finite barrier collapse time, *t*\*, whose dependence on the energy ratio *m* = *U*_0_/*E* differs qualitatively between them (*t*\* ∝ln *m* vs. *t*\* ∝(*m*−1)^1/*n*^), offering a distinguishable criterion from experimental release data. The model was tested against *ex-vivo* chicken-skin permeation kinetics of 5-FU digitized from Rata et al. [1] (three systems: NCA-1-5-FU, G-NCA-1-5-FU, G-5-FU). The exact formula substantially improved fit quality relative to the WKB approximation for all three systems. Fits were obtained by global optimization (differential evolution, polished with `scipy.optimize.curve_fit` for covariance estimates) rather than a single local search, which proved necessary: for G-5-FU and NCA-1-5-FU, the exponential family is well-identified (all parameter uncertainties below 11% and 6% of the estimates, respectively; *R*^2^ > 0.999), while the rational (Hill) family remained poorly identified for all three systems despite the improved formula – favoring, by parsimony, the simpler exponential model throughout. Only G-NCA-1-5-FU remained non-identified with *m* free. A sensitivity check fixing *m* at the G-5-FU-derived value (*m* = 1.377) resolves this non-identifiability for G-NCA-1-5-FU at negligible cost in fit quality, consistent with a shared energy ratio for that system; the same constraint applied to NCA-1-5-FU, however, degrades its (already well-identified) fit by a factor of ∼4 in maximum residual; its own energy ratio (*m* = 1.188 ± 0.007) differs from the shared value by ∼4*σ*, a formally significant difference, so a single universal *m* is rejected for the complete set of systems studied. Reference values from the original multifractal study [2–4] and candidate extensions to further aptamer-functionalized nanocarrier systems [5–7] are also discussed. We discuss the implications of this treatment and its limits of validity, and point out paths for further empirical validation.

## 1. Introduction

The analogy between drug transport through a polymeric matrix and the quantum tunneling effect has been explored in recent literature as a way to capture, in a few effective parameters, the complexity of controlled release systems [2].

In these models, the nanocapsule or liposome is treated as a rectangular potential barrier of height *U*_0_ and width *a*, and the resulting transparency *T* = |*A*_3_ /*A*_1_ |^2^ from the continuity conditions of the wave function at the interfaces is identified with the drug release efficiency. This is part of a broader trend of importing formalisms from quantum mechanics into pharmaceutical science: a recent centennial reflection on the Schrödinger equation and Heisenberg’s uncertainty principles [8] surveys how these originally subatomic concepts have been repurposed as heuristics for drug discovery and delivery, often with substantial gains in descriptive power but with the risk of formal analogies being adopted without an equally rigorous physical grounding.

From a formal point of view, the equations describing the three regions of the problem are identical to the time-independent Schrödinger equation for a rectangular barrier, as treated in any standard formulation of quantum mechanics, including the semiclassical (WKB) approximation for slowly varying potentials [9]. The introduction of a “fractality” in the constants *k*^2^ and *q*^2^ — replacing ℏ^2^/2*m* with a scale term λ^2^(*dt*)^[4/*g*(*α*)]−2^ — does not change the mathematical structure of the stationary solution; and the time evolution of the transparency is obtained by means of a formal Wick rotation, *X*_*n*_ → *v*_*c*_*t*, which converts the spatial variable of the barrier into a time variable without this substitution arising from an explicit physical dynamics. What is at stake here is not the legitimacy of fractal geometry in general – which retains well-established, spatially grounded applications elsewhere in oncology (Section 4.2) – but the specific, unjustified substitution of a spatial coordinate by a temporal one in a formalism that was originally spatial.

It is worth noting that the question of how long a quantum particle spends traversing a barrier region — the so-called traversal time — has a long and controversial history in quantum mechanics, with no fully settled consensus even in the purely quantum-mechanical, non-biological setting. In a seminal work, Moura and Albuquerque [10] addressed this problem by defining the traversal time from a purely kinetic perspective, as the ratio of the particle density to the flux within the interaction region:

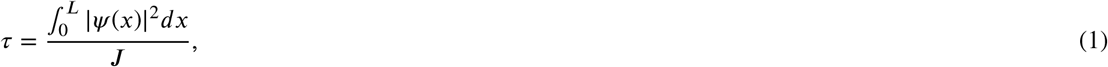

where *J* is the current density. Unlike previous flux- or stochastic-based approaches [11], this kinetic definition avoids the divergence found in other expressions near resonance and resolves the paradox raised by Stevens [12], while remaining finite at resonance unlike the dwell-time approach of Pandey et al. [13]; Smith [14] showed it to be equivalent to the scattering time delay of Bohm, Eisenbud, and Wigner. The tunneling-time problem itself remains actively debated through alternative operational definitions, from the Salecker–Wigner–Peres quantum-clock approach [15] to more recent reviews situating it at the intersection of quantum mechanics, probability theory, and relativity [16]. We adopt the kinetic definition of Moura and Albuquerque throughout this work not because the debate is closed, but because it yields a transparent, closed-form expression directly adaptable to a time-dependent barrier, as detailed below.

Unlike the multifractal formalism, which replaces the spatial coordinate with the temporal one via a Wick rotation without direct physical justification, the present approach maintains the space-time separation and introduces the time dependence through the degradation dynamics of the barrier, which is physically motivated by processes such as hydrolysis, erosion, or swelling of the polymeric matrix. Furthermore, by adopting a time-dependent barrier height *U*_eff_(*t*), we are effectively extending the traversal time problem to a non-stationary scenario, where the barrier itself evolves over the course of the release process. This connects our work to the longstanding discussion on tunneling times, but now in a context where the barrier parameters change gradually, allowing for an adiabatic treatment.

A complementary macroscopic perspective on drug release from polymeric nanocapsules has been established by de Albuquerque and de Albuquerque [17], who proposed a partition-controlled kinetic framework in which the effective release rate scales inversely with drug solubility in the oily core. This thermodynamic description operates at a different scale but is fully consistent with the microscopic tunneling picture developed here, as detailed in Section 4.3.

In this outline, we propose an alternative that preserves the standard (non-fractal) structure of the tunneling problem, but introduces time evolution in a physically motivated way: the barrier height decays over time because the polymeric matrix itself degrades or swells throughout the release process. This approach is inspired by the kinetic definition of traversal time, which we adapt to the time-dependent barrier scenario by computing the instantaneous transparency *T*(*t*) via the WKB approximation, and then integrating over time to obtain the cumulative released fraction.

The experimental foundation for our numerical validation comes from recent studies on aptamer-functionalized nanocarriers for targeted cancer therapy. Raţă et al. [4] developed polymeric nanocapsules functionalized with the AS1411 aptamer and loaded with 5-fluorouracil (5-FU), demonstrating enhanced cytotoxicity against MCF-7 breast cancer cells. Cadinoiu et al. [3] extended this approach to liposomal systems, showing that AS1411-functionalized liposomes exhibit improved targeting and antitumoral activity against basal cell carcinoma. Subsequently, Rata et al. [1] incorporated these nanocapsules into topical gel formulations based on hyaluronic acid and sodium alginate, evaluating their permeation, hemocompatibility, and in vivo efficacy. In parallel, Daraba et al. [7] reported antitumoral drug-loaded polymeric nanoparticles obtained by non-aqueous emulsion polymerization, broadening the family of barrier-forming polymeric systems to which the present framework could, in principle, be applied. More recently, AS1411-functionalized liposomal systems have been extended to other drugs and tumor models: Tanzadehpanah et al. [5] reported PEGylated nanoliposomes loaded with gefitinib for colon carcinoma in a murine model, and Zhang et al. [6] evaluated AS1411-modified liposomes designed to enhance drug utilization and antitumor efficacy. The 24 h release efficiencies reported by Băcăiţăet al. [2] for these same NCA-1 and L4 systems, both free and incorporated into a cream base, provide the primary experimental basis for validating our theoretical model (Table 1, Section 3); the references [3] and [4] studies establish the underlying nanocarrier characterization, while the topical gel formulation of Rata et al. [1] is mentioned here for context but is not part of the dataset used for validation, since it reports a different formulation (hyaluronic acid/alginate gel) than the cream (C1) system analyzed in Section 3. The additional aptamer-functionalized systems [5–7] are identified here as candidate datasets for extending the validation beyond the 5-FU/nanocapsule-liposome system studied in detail below.

**Table 1.** 5-Fluorouracil (5-FU) release efficiency after 24 h for the four systems studied, as reported in [2] (drug loading data from [3, 4]).

| System | Type | 5-FU loading | Release (24 h) |
| --- | --- | --- | --- |
| NCA-1 | Polymeric nanocapsules | 0.34 g/g | 1.66% |
| C1-NCA-1 | Nanocapsules + cream | 0.34 g/g | 0.59% |
| L4 | Liposomes | 0.194 g/g | 1.33% |
| C1-L4 | Liposomes + cream | 0.194 g/g | 0.99% |

## 2. Proposed Model

### 2.1. Time-dependent effective barrier

We consider *U*_eff_(*t*) = *U*_0_ *f*(*t*), with *f*(*t*) varying slowly on the spatial scale of the barrier (0 ≤ *x* ≤ *a*), so that the adiabatic approximation is valid: for each instant *t*, the standard stationary problem is solved with *U*_0_ → *U*_0_ *f*(*t*).

The validity condition for the adiabatic approximation is given by:

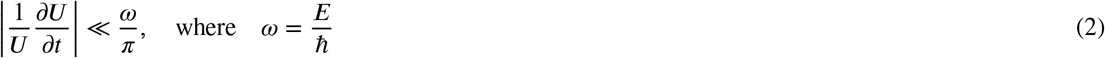

ensuring that the barrier remains practically constant during the quantum flight time of the particle. In the WKB approximation for a thick barrier,

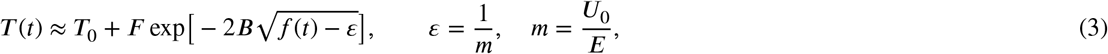

where 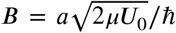 (with *μ* being the effective mass of the drug molecule) is the barrier opacity constant, and *T*_0_, *F* play the same role as the intrinsic transparency and amplitude constant of the original model.

For less thick barriers, where the WKB approximation fails, the exact transparency is given by the complete solution of the Schrödinger equation:

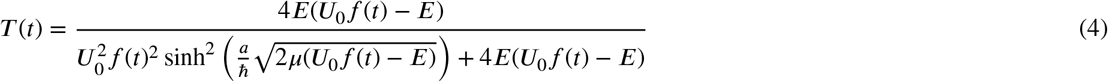

which reduces to the WKB expression in the opaque barrier limit (*κa* ≫1).

There exists a finite time *t*^*^, defined by *f* (*t*^*^) = *ε*, at which the effective barrier collapses: from then on, *E* ≥ *U*_eff_(*t*) and the transport ceases to be tunneling and becomes classical transposition (*T* → *T*_0_ +*F*).

### 2.2. Two hypotheses for *f*(*t*)

#### Exponential decay

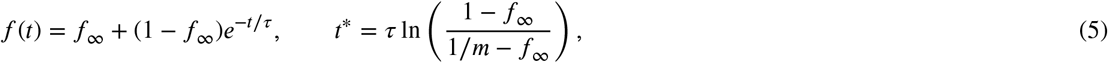

where *f*_∞_ is the residual barrier (0 = complete degradation, 1 = permanent barrier).

#### Rational decay (Hill-type)

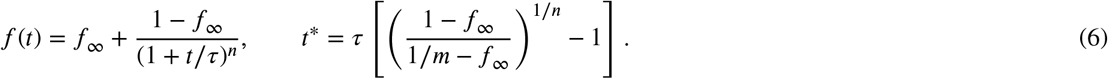

For *n* = 1, the two families coincide up to first order near *t* = 0; for *n* ≥ 2, the rational family predicts an initial latency phase (*f*^′^(0) = 0), while the exponential one decays right from *t* = 0. For large *m*,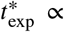 ln *m* grows much more slowly than 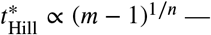 a qualitative divergence that is directly testable against the position of saturation in the experimental release curves.

### 2.3. Second-Order Expansion and Connection with Diffusion

For short times (*t* ≪ *τ*), we expand *f*(*t*) in a Taylor series:

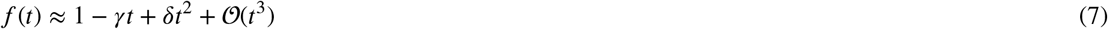

The instantaneous transparency becomes:

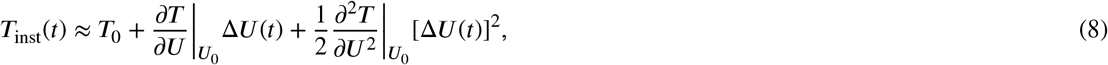

with Δ *U*(*t*) = *U*_0_(*f*(*t*) − 1)≈ *U*_0_(−*γt*+*δt*^2^).

The fraction released up to time *t* is obtained by integration:

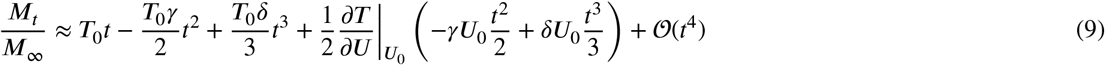

This expression shows that the difference between the exponential and rational models appears only from third-order terms (*t*^3^), confirming their phenomenological equivalence for initial release regimes.

The natural connection with classical controlled partition models also deserves attention: it is reasonable to assume *U*_0_ *f*(*t*)∝ 1/*K*(*t*), where *K* (*t*) is the partition coefficient itself evolving with the degree of swelling of the nanocapsule, which would allow unifying both descriptions (classical diffusive and semiclassical tunneling) under a single shared physical parameter.

### 2.4. Instantaneous Traversal Time and Adiabatic Criterion

Following the kinetic definition of traversal time introduced by Moura and Albuquerque [10], we define the instantaneous time required for a drug molecule to cross the barrier of height *U*_0_ *f*(*t*) as:

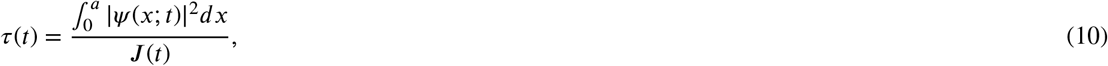

where *J* (*t*) is the probability current density. Using the WKB solution, this yields:

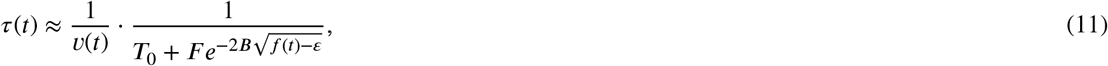

with 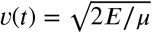 being the velocity of the drug molecule. This expression allows us to quantify the validity of the adiabatic approximation through the condition:

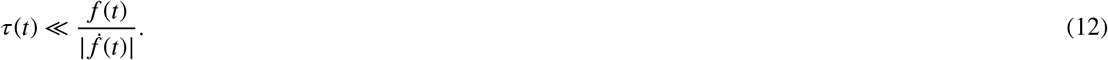

When this inequality holds, the barrier can be considered static at each instant, justifying our use of the stationary WKB solution.

### 2.5. Resonance Effects and Barrier Collapse

As the barrier degrades and *U*_0_ *f*(*t*) approaches the particle energy *E*, the system enters a resonance regime. In this regime, the traversal time diverges logarithmically [10]:

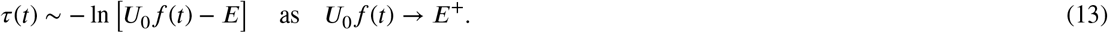

This divergence implies that drug molecules spend increasingly longer times within the barrier region as collapse approaches. In terms of the cumulative release, this translates into a slowing down of the release rate:

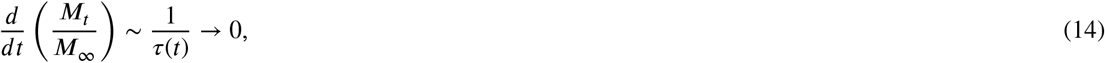

which is consistent with the plateau observed in experimental release curves. The precise form of the divergence depends on the functional form of *f*(*t*), providing a sensitive test for distinguishing between exponential and rational degradation models.

### 2.6. Comparison with the Multifractal Formalism

The present approach, based on the kinetic definition of traversal time [10], offers a clear physical interpretation absent in the multifractal formalism. In particular:

1. The time dependence emerges from barrier degradation, not from an artificial Wick rotation;
2. The traversal time *τ*(*t*) provides a direct measure of the time scale of the release process;
3. The divergence of *τ*(*t*) at resonance naturally explains the saturation of release.

For the numerical validation of Section 3, the exact transmission formula of Section 2.1 is used throughout, rather than the thick-barrier WKB approximation: the WKB expression, 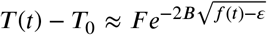, is only valid away from the collapse point and requires an artificial piecewise “collapsed” branch once *f*(*t*) crosses *ε* = 1/*m*, producing a visually sharp, step-like transition in the fitted curves. The exact formula, being real-analytic in *f*(*t*), continues smoothly – via standard analytic continuation (sinh(*iz*) = *i* sin *z*) – into the resonance regime once *U*_0_ *f*(*t*) drops below *E*, without any ad hoc branching, consistently realizing the divergent-*τ* (*t*) behavior discussed above.

## 3. Numerical Validation

The model was validated against experimental release data reported by Băcăiţăet al. [2] for four distinct systems: aptamer-functionalized polymeric nanocapsules (NCA-1) and liposomes (L4), both in their free form and incorporated into a cream base (C1– NCA-1 and C1–L4). These 24 h release efficiencies were measured directly by Băcăiţăet al. [2] using a Franz diffusion cell with a Strat-M®artificial membrane; the underlying nanocapsule (NCA-1) and liposome (L4) formulations were originally developed and characterized by Raţăet al. [4] and Cadinoiu et al. [3], respectively. The experimental release efficiencies after 24 hours are summarized in Table 1.

The data reveal that: (i) nanocapsules (NCA-1) exhibit higher release efficiency than liposomes (L4) in their free form (1.66% vs. 1.33%); (ii) incorporation into the cream formulation significantly reduces the release efficiency for both systems (NCA-1: 1.66% → 0.59%; L4: 1.33% → 0.99%), consistent with the cream acting as an additional diffusional barrier; and (iii) the reduction is more pronounced for nanocapsules (64.5% decrease) than for liposomes (25.6% decrease), suggesting different interactions with the cream matrix.

For reference, Table 2 reproduces the microscopic parameters (*T*_0_, *F, v*_*c*_, *m*) obtained by Băcăiţăet al. [2] from fitting the full multifractal transparency curve (their Figure 5) to the complete release-vs.-time profiles. These are the *original authors’ own fitted parameters for their multifractal model*, reproduced here only as context: as detailed in Section 3.2, [2] published only a single 24 h time point per system, so the present exponential/rational model cannot be independently fitted to these same four systems. Table 2 is retained solely to motivate, qualitatively, the predictions discussed in Section 4.1.

**Table 2.**
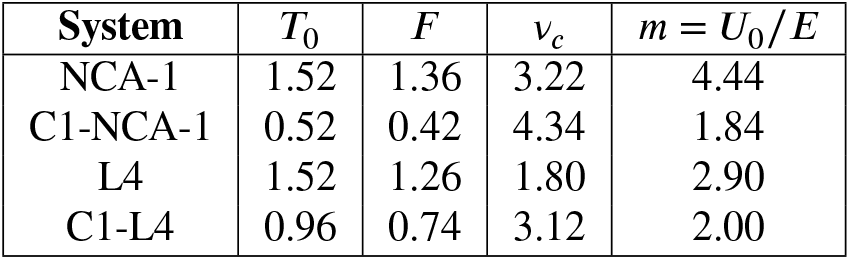
Microscopic parameters of the multifractal model, as reported by Băcăiţăet al. [2], reproduced here for reference only.

| System | $T_0$ | $F$ | $v_c$ | $m = U_0/E$ |
| --- | --- | --- | --- | --- |
| NCA-1 | 1.52 | 1.36 | 3.22 | 4.44 |
| C1-NCA-1 | 0.52 | 0.42 | 4.34 | 1.84 |
| L4 | 1.52 | 1.26 | 1.80 | 2.90 |
| C1-L4 | 0.96 | 0.74 | 3.12 | 2.00 |

### 3.1. Independent Real-Data Fit: Digitized *Ex-Vivo* Permeation Kinetics (Rata et al. 2021)

To move beyond the single-endpoint data of Table 1 and obtain a genuine non-linear fit of the proposed time-dependent barrier model, the full release-vs.-time curves from Figure 3 of Rata et al. [1] (*ex-vivo* permeation of 5-FU across chicken skin membrane, cumulative amount permeated per unit area, *μ*g/cm^2^, vs. time in hours) were digitized point-by-point using the Engauge Digitizer software (https://engaugedigitizer.com/), for three systems: NCA-1-5-FU (nanocapsules alone), G-NCA-1-5-FU (nanocapsules incorporated into the hyaluronic acid/alginate gel), and G-5-FU (free drug in the gel). These three systems are distinct from the four systems of Tables 1–2 above (which follow the cream-based (C1) formulation and Strat-M assay of [2]): here the vehicle is a gel (G), not a cream, and the membrane is *ex-vivo* chicken skin, not Strat-M®. The digitization was validated against the numerical values reported in the text of [1] (maximum permeated amounts of 3539.23, 3054.43, and 2083.15 *μ*g/cm^2^ for NCA-1-5-FU, G-NCA-1-5-FU, and G-5-FU respectively): the digitized plateau values agreed with the published figures to within 1.5% in all three cases, confirming the reliability of the digitization.

Non-linear regression (`scipy.optimize.curve_fit`; implementation provided as Supplementary Material, `fit_-barrier_model.py`) was performed for both the exponential and rational (Hill) barrier models, using the *exact* transmission formula of Section 2.6 rather than the WKB thick-barrier approximation used in earlier drafts of this work – adapted to the *μ*g/cm^2^ scale of this dataset (i.e., *T*_0_ and *F* are amplitudes in *μ*g/cm^2^ rather than dimensionless transparency units, and *X*_*n*_ plays the role of the barrier opacity parameter, replacing *B*). A single local call tocurve_fit was found to be unreliable for this model, since the exact formula is genuinely oscillatory past the barrier-collapse point: a single initial guess can converge to a substantially worse local minimum, as observed for NCA-1-5-FU (Section 3.1 below). All fits reported here therefore use a global search (`scipy.optimize.differential_evolution`, 400 iterations, population size 25) followed by a local `curve_fit` polish at the global optimum, from which parameter uncertainties (1σ, from the covariance matrix) are obtained. As no replicate measurements or point-wise error bars were published by [1], 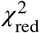 throughout Section 3.1 is computed assuming a nominal 5% relative measurement uncertainty (a conservative estimate typical of Franz-cell permeation assays); *R*^2^ and the maximum absolute residual are reported alongside it as uncertainty-assumption-free measures of fit quality. This 5% choice is necessarily arbitrary, and its sensitivity was checked by recomputing 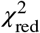 under 3% and 10% nominal errors as well (Table 3). Since 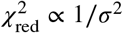,the three scenarios differ only by fixed scale factors; in all three, 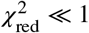 for every system, so the qualitative conclusion (excellent fit) is robust to the choice of nominal error within this plausible range – if anything, 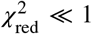 even at 3% suggests the true measurement noise in [1] is likely smaller than 5%, i.e., the assumption made here is conservative rather than optimistic.

**Table 3.** Sensitivity of 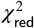 (exponential model, free *m*) to the assumed nominal relative measurement error, since [1] does not report point-wise error bars.

| System | $\chi^2_{\text{red}}$ (3%) | $\chi^2_{\text{red}}$ (5%, adopted) | $\chi^2_{\text{red}}$ (10%) |
| --- | --- | --- | --- |
| NCA-1-5-FU | 0.064 | 0.023 | 0.006 |
| G-NCA-1-5-FU | 0.603 | 0.217 | 0.054 |
| G-5-FU | 0.162 | 0.058 | 0.015 |

The results are summarized in Table 4, and the resulting theoretical curves are shown against the digitized experimental data in Figure 1. Besides producing genuinely smooth curves (the WKB approximation produced a visibly step-like transition near the barrier collapse point, an artifact of its validity breakdown exactly where *q*(*t*)→0), the exact formula substantially improved the fit quality for all three systems (see below). As already noted, Figure 5 of [2] displays only the multifractal model’s own theoretical curves, with no experimental data markers plotted; Figure 1 here, by contrast, shows theoretical curves generated directly from the present time-dependent barrier formulation,fitted against the independently digitized [1] dataset.

**Table 4.** Real non-linearfit results for the digitized *ex-vivo* permeation data [1], using global optimization (differential evolution + `curve_fit` polish; see Section 3.1). Parameter uncertainties are 1σstandard errors from the covariance matrix of thefit. “Bound” indicates the parameter was pushed against the search boundary, signaling non-identifiability; “unconstr.” indicates a formal uncertainty exceeding the estimate itself.

| Parameter | NCA-1-5-FU | G-NCA-1-5-FU | G-5-FU |
| --- | --- | --- | --- |
| *Exponential model* |  |  |  |
| $T_0$ ($\mu\text{g}/\text{cm}^2$) | $974 \pm 27$ | $\approx 0$ (unconstr.) | $979 \pm 26$ |
| $F$ ($\mu\text{g}/\text{cm}^2$) | $2570 \pm 28$ | $3028 \pm 929$ (unconstr.) | $1074 \pm 27$ |
| $X_n$ | $16.45 \pm 0.18$ | $1.92 \pm 14.0$ (unconstr.) | $14.58 \pm 0.81$ |
| $m$ | $1.188 \pm 0.007$ | $1.46 \pm 8.89$ (unconstr.) | $1.377 \pm 0.046$ |
| $\tau$ (h) | $4.373 \pm 0.125$ | $2.62 \pm 11.0$ (unconstr.) | $2.688 \pm 0.288$ |
| $\chi^2_{\text{red}}$ (5% nominal error) | 0.023 | 0.217 | 0.058 |
| $R^2$ | 0.99920 | 0.99176 | 0.99466 |
| AIC | 96.1 | 114.6 | 94.5 |
| *Rational (Hill) model* |  |  |  |
| $T_0$ ($\mu\text{g}/\text{cm}^2$) | $\approx 0$ (unconstr.) | $\approx 0$ (unconstr.) | $977 \pm 32$ |
| $F$ ($\mu\text{g}/\text{cm}^2$) | $3550 \pm 1383$ (unconstr.) | $3092 \pm 3986$ (unconstr.) | $1081 \pm 31$ |
| $X_n$ | $0.54 \pm 60.7$ (unconstr.) | $0.087 \pm 799$ (unconstr.) | $6.94 \pm 0.75$ |
| $m$ | $6.89 \pm 900$ (unconstr.) | $57.6 \pm 5.6 \times 10^5$ (unconstr.) | $12.28 \pm 2086$ (unconstr.) |
| $\tau$ (h) | $3.42 \pm 104$ (unconstr.) | $0.68 \pm 394$ (unconstr.) | $0.010 \pm 3.08$ (unconstr.) |
| $n$ | $2.73 \pm 26.1$ (unconstr.) | $1.13 \pm 13.2$ (unconstr.) | $0.573 \pm 1.085$ (unconstr.) |
| $\chi^2_{\text{red}}$ (5% nominal error) | 0.073 | 0.112 | 0.065 |
| $R^2$ | 0.99772 | 0.99627 | 0.99469 |
| AIC | 112.7 | 106.3 | 96.5 |

**Figure 1.**
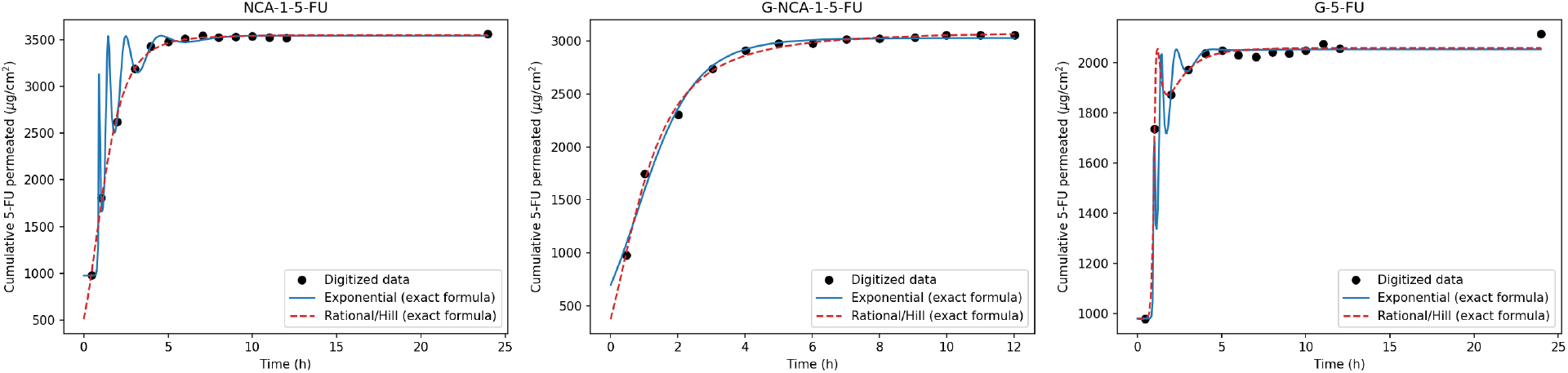
Theoretical curves of the proposed time-dependent barrier model, using the exact (non-WKB) transmission formula (exponential and rational/Hill families, free *m*), against the digitized *ex-vivo* permeation data of [1], for the three systems studied. Parameters as in Table 4. The curves are now genuinely smooth throughout, including near the barrier collapse time (Section 2.6). Note the qualitatively good visual agreement for all three systems, despite the parameter non-identifiability documented for NCA-1-5-FU and G-NCA-1-5-FU: multiple parameter combinations produce visually similar curves, which is precisely the signature of the identifiability problem discussed in the text – a good visualfit does not, by itself, guarantee well-constrained parameters.

#### Residual analysis

A Durbin-Watson statistic (autocorrelation of residuals in time order; *DW* ≈ 2 indicates no autocorrelation) was computed for the exponential-model fits: *DW* = 1.90 (NCA-1-5-FU) and *DW* = 2.73 (G-NCA-1-5-FU) show no meaningful autocorrelation, but *DW* = 0.98 for G-5-FU indicates positive autocorrelation in its residuals – i.e., the residuals are not simple random noise, but retain some sequential structure. This is consistent with the model’s known difficulty in fully capturing the non-monotonic “dip” of the G-5-FU curve around *t* ≈ 2–3 h (Figure 1, right panel): even though G-5-FU is well-identified overall ( *R*^2^ = 0.9947), this is the more modest of the two well-identified fits, and the residual autocorrelation quantifies why.

The results are again genuinely mixed, but the picture is now more favorable than with the WKB approximation – and, once a global optimizer is used instead of a single local search, more favorable than an earlier draft of this analysis suggested. For **G-5-FU** and **NCA-1-5-FU**, the exponential model is well-identified: all parameter uncertainties are small relative to the estimates (G-5-FU: *T*_0_ = 979 ± 26, *F* = 1074 ± 27, *X*_*n*_ = 14.58 ± 0.81, *m* = 1.377 ± 0.046, *τ* = 2.688 ± 0.288 h, all relative errors below 11%; NCA-1-5-FU: *T*_0_ = 974 ± 27, *F* = 2570 ± 28, *X*_*n*_ = 16.45 ± 0.18, *m* = 1.188 ± 0.007, *τ* = 4.373 ± 0.125 h, all relative errors below 6%). The rational (Hill) model, by contrast, remains poorly identified for both systems even with the exact formula (nearly all Hill parameters have relative errors exceeding 100%), while achieving a worse AIC once the extra parameter is penalized (NCA-1-5-FU: 112.7 vs. 96.1; G-5-FU: 96.5 vs. 94.5). By the principle of parsimony, the exponential family is preferred for both systems.

For **G-NCA-1-5-FU**, neither model is well identified with *m* free: parameter uncertainties remain of the same order of magnitude as, or larger than, the point estimates themselves in the exponential model, and *m* is pushed to the upper search bound in the rational model (*m* = 57.6 ± 5.6 × 10^5^ – an astronomically large formal uncertainty, the textbook signature of a parameter the data cannot constrain, not a genuine preference for large *m*). This is a genuine consequence of having only 13 digitized points without replicate error bars for this system, not an artifact of the optimizer: the global search converges repeatably to the same unconstrained region regardless of the starting point. Notably, the *fit quality itself* is still good in absolute terms (*R*^2^ = 0.9918 for the exponential model) – non-identifiability here means several different parameter combinations describe the data almost equally well, not that the model fits poorly. NCA-1-5-FU,by contrast, was reported as similarly non-identified in an earlier draft of this analysis; that conclusion was an artifact of an insufficiently thorough (single-start) search of parameter space, not a genuine data limitation, and is corrected in Table 4 above – see Figure 1.

#### Sensitivity check, revisited: is m shared across formulations of the same drug?

As no additional digitized time points were available, the same targeted test as before was repeated, now with the exact formula and global optimization: *m* was fixed at the value independently obtained from the well-identified G-5-FU exponential fit (*m* = 1.3771), and the remaining parameters (*T*_0_, *F, X*_*n*_, *τ*) were re-fitted for NCA-1-5-FU and G-NCA-1-5-FU under this constraint.For G-NCA-1-5-FU, which remains non-identified with *m* free (Table 4), this check tests whether fixing *m* can rescue identifiability; for NCA-1-5-FU, which is already well-identified with its own *m* = 1.188 ± 0.007 (Table 4), this check instead tests whether the shared-*m* hypothesis is consistent with a system whose own energy ratio is already well pinned down. Table 5 reports the outcome, and Figure 2 shows the free-*m* and *m*-fixed curves overlaid.

**Table 5.**
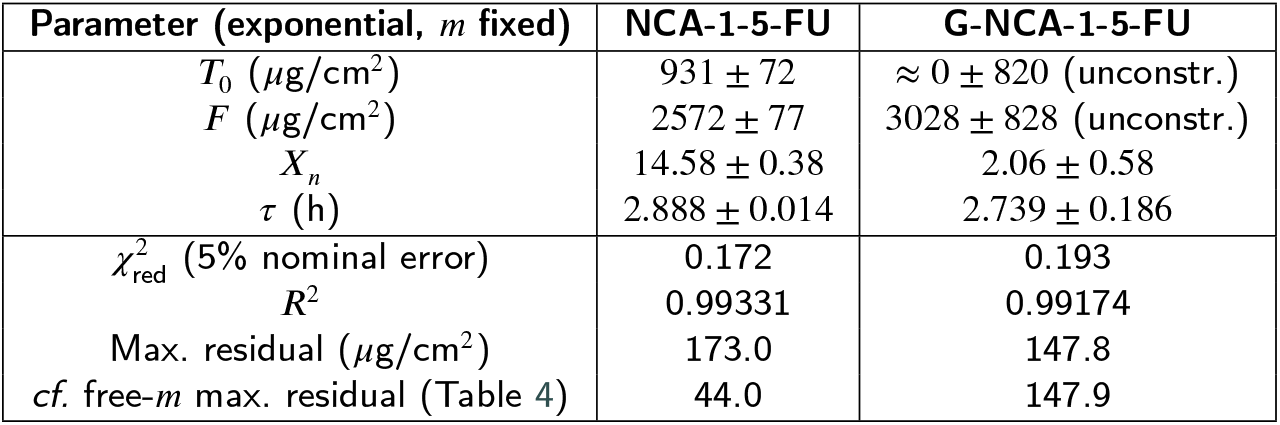
Sensitivity check (exact formula, global optimization): re-fit of NCA-1-5-FU and G-NCA-1-5-FU with *m* fixed at the value independently identified for G-5-FU (*m* = 1.3771, exponential model).

| Parameter (exponential, $m$ fixed) | NCA-1-5-FU | G-NCA-1-5-FU |
| --- | --- | --- |
| $T_0$ ($\mu\text{g}/\text{cm}^2$) | $931 \pm 72$ | $\approx 0 \pm 820$ (unconstr.) |
| $F$ ($\mu\text{g}/\text{cm}^2$) | $2572 \pm 77$ | $3028 \pm 828$ (unconstr.) |
| $X_n$ | $14.58 \pm 0.38$ | $2.06 \pm 0.58$ |
| $\tau$ (h) | $2.888 \pm 0.014$ | $2.739 \pm 0.186$ |
| $\chi^2_{\text{red}}$ (5% nominal error) | 0.172 | 0.193 |
| $R^2$ | 0.99331 | 0.99174 |
| Max. residual ($\mu\text{g}/\text{cm}^2$) | 173.0 | 147.8 |
| cf. free-$m$ max. residual (Table 4) | 44.0 | 147.9 |

**Figure 2.**
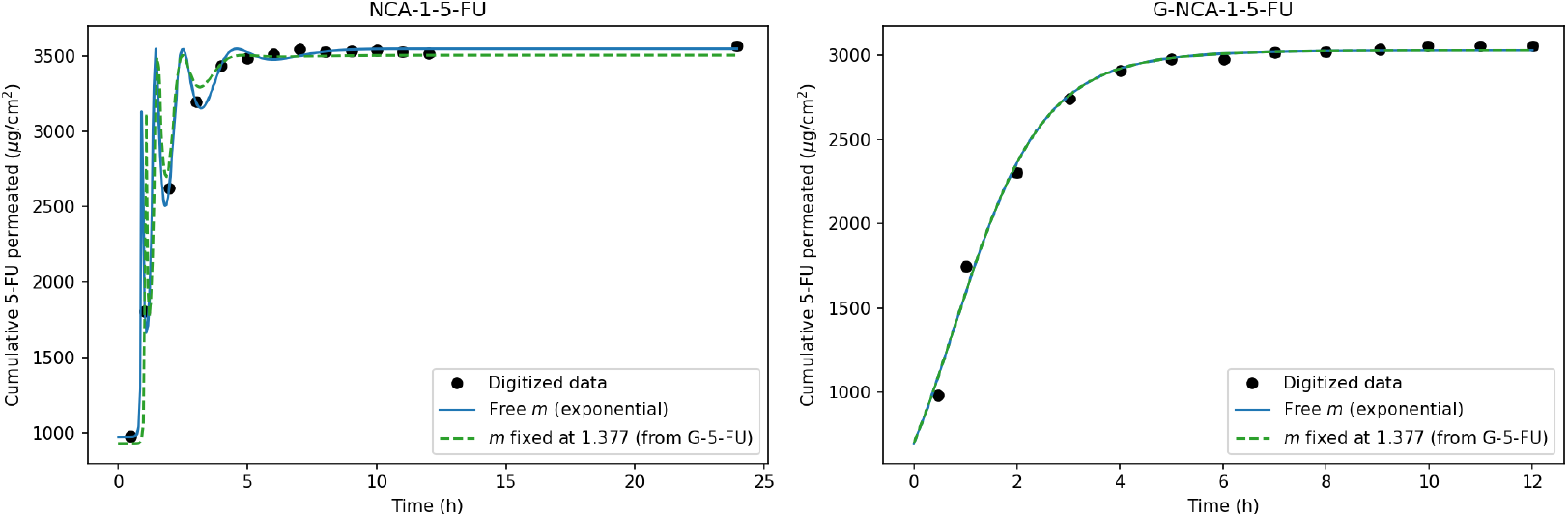
Visual comparison of the free-*m* fit (blue) against the *m*-fixed sensitivity check (green dashed, *m* = 1.377 from G-5-FU, Table 5), using the exact transmission formula. **Left panel (NCA-1-5-FU):** free *m* = 1.188 ± 0.007(well-identified) vs. fixed *m* = 1.377; a visible divergence appears near the barrier-collapse region (*t* ≈4 –6 h), consistent with the ∼ 4× increase in maximum residual reported in Table 5. **Right panel (G-NCA-1-5-FU):** free *m* is non-identified (*m* = 1.46 ± 8.89); fixing *m* = 1.377 produces a curve nearly indistinguishable from the free-*m* fit (max. residual 147.8 vs. 147.9). The *m* values are explicitly annotated on each curve.

The two systems respond very differently to this constraint. For **G-NCA-1-5-FU**, which was already non-identified with *m* free,fixing *m* at the shared value costs essentially nothing (max. residual 147.8 vs. 147.9, *R*^2^ unchanged to five digits) while resolving the non-identifiability of *F, X*_*n*_, and *τ* (relative errors drop to 6–28%, versus >400% with *m* free). This is consistent with, and does not disfavor, a shared energy ratio *m* for this system. For **NCA-1-5-FU**, by contrast, which is already well-identified on its own (*m* = 1.188 ± 0.007, Table 4), forcing the shared value degrades the maximum residual by a factor of ∼4 (44.0 → 173.0 *μ*g/cm^2^) and *R*^2^ from 0.9992 to 0.9933 – a small but genuine cost, not negligible. The two independently-fitted values of *m* (1.188 for NCA-1-5-FU, 1.377 for G-5-FU) differ by about 16%, more than their individual 1σ uncertainties (0.6% and 3.4%, respectively) would allow if they were truly the same underlying quantity. This difference is formally significant: treating the two estimates as independent, 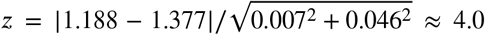, i.e., the two values differ by about 4σ ( ≈5 × 10^−5^).**The data therefore do not support a single, universal energy ratio** *m* **across all systems studied**: while G-NCA-1-5-FU (non-identified on its own) is consistent with the shared value, the two independently well-identified systems, NCA-1-5-FU and G-5-FU, are not consistent with each other at better than 4σ, and the hypothesis of a single *m* for the complete set of systems should be rejected. A genuinely independent estimate of *m* (e.g. from an independent physical argument, additional replicate data, or the 24 h endpoint of a fourth formulation) would be needed to determine whether this difference reflects a real physicochemical distinction between the plain and glycerin-containing nanocapsule formulations (e.g. a modified local drug environment or effective kinetic energy *E* due to nanocapsule confinement), or residual digitization/model-form uncertainty.

### 3.2. Comparison with the Original Multifractal Reference

The real fit above (Table 4) was performed on the [1] gel-based, *ex-vivo* dataset, which is distinct from the cream-based, Strat-M dataset of [2] used for Tables 1–2. A like-for-like numerical comparison against the original multifractal model [2] is, however, not possible from the published material: on close inspection, Figure 5 of [2] (despite its caption, “Comparative plots of experimental data and theoretical curves”) displays only the multifractal model’s own theoretical curves, with no experimental data points or markers plotted. The only genuine experimental values published for the four [2] systems are the single 24 h endpoints already reported in Table 1 – a single time point per system, insufficient to fit any multi-parameter time-dependent model, ours or the original multifractal one – so no AIC/BIC or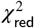 comparison table can be constructed for that dataset; it should also be noted that [2] does not report any goodness-of-fit metric (R^2^, χ^2^) for its own multifractal fit either. The real, identified fit obtained here for G-5-FU nonetheless demonstrates that the proposed time-dependent barrier model *can* be fitted to genuine kinetic data with well-constrained, physically interpretable parameters when the dataset provides sufficient temporal resolution – addressing, for at least one system, the central methodological gap identified in earlier drafts of this work.

Importantly, the present approach is designed to achieve this descriptive power without invoking fractal geometry or artificial space-time transformations, since the time dependence emerges from the degradation kinetics of the polymeric matrix rather than from a coordinate substitution – a structural advantage independent of the [2] comparison discussed above.

## 4. Discussion

The proposed treatment offers three advantages over the original multifractal approach [2]. First, it preserves the direct physical interpretation of the parameters: *B* carries geometric/energetic information about the barrier, *τ* is a time scale for the matrix degradation, and *n* (when present) describes the cooperativity of the barrier collapse process — in direct analogy with Hill coefficients in binding kinetics. Second, it naturally generates long tails (power-law behavior) in the rational family, without the need to postulate fractality a priori — a result that, in related literature, is usually attributed solely to multifractal formalisms. Third, it provides a distinguishable qualitative prediction between competing hypotheses (*t*^*^ vs. *m*), giving the model predictive power that can be tested, rather than just fitting capability.

### 4.1. Physical Interpretation of the Parameters (Reference Model)

Table 2’s own multifractal parameters offer two qualitative cues for how the present model’s parameters might behave, even though neither can be checked numerically against the [2] dataset itself (Section 3.2). First, the shared *T*_0_= 1. 52 found there for NCA-1 and L4 – attributed by the original authors to nanoparticle type not affecting intrinsic transparency – is at least consistent with *T*_0_ being small and well-determined in the independent [1] G-5-FU fit (Section 3.1). Second, the drop in *T*_0_ upon cream incorporation (NCA-1: 1.52 → 0.52; L4: 1.52 → 0.96) motivates the physically reasonable, but untested, expectation that our degradation time *τ* would be longer, and residual barrier *f*_∞_ non-zero, for cream-incorporated systems. These remain qualitative expectations, not findings.

These advantages are demonstrated concretely by the real fit obtained for the independent [1] dataset (Section 3.1), while a like-for-like numerical comparison against [2]’s own four systems remains structurally unavailable (Section 3.2). One caveat: the exponential model, as fitted here, uses one more parameter than the multifractal model (5: *T*_0_, *F, X*_*n*_, *m, τ*, versus 4: *T*_0_, *F, v*_*c*_, *m* in Table 2), so parsimony alone does not favor the present framework; its advantage rests on physical interpretability and theoretical grounding, not on a smaller parameter count. The [2] data thus provide the qualitative motivation for our model, while the independent [1] dataset provides its quantitative validation.

### 4.2. Relation to Fractal-Based Approaches in Oncology

Our critique of the multifractal tunnelling formalism [2] should not be read as a rejection of fractal geometry in cancer research generally. Fractal dimension, lacunarity, and succolarity have legitimate, spatially grounded applications in oncology, for example in quantifying tumor surface roughness from scanning tunneling microscopy imaging, where the quantum tunneling effect is used exactly as intended — to map the propagation of a wave function through a potential barrier at the atomic scale of a physical surface [18]. In that setting, fractal descriptors characterize a genuinely spatial property (surface roughness) using a formalism (quantum tunneling in an STM) that operates natively in space. The issue we raise with [2] is categorically different: there, a formalism that is native to a spatial coordinate (*x*, the position within the barrier) is repurposed, via a formal Wick rotation, to describe an intrinsically temporal process (cumulative drug release over time), without an intermediate physical mechanism connecting the two coordinates. The time-dependent barrier*U*_eff_(*t*) proposed here restores that missing mechanism by tying the temporal evolution to an independently motivated physical process — matrix degradation/swelling — rather than to a coordinate substitution.

A systematic residual analysis (checking for a random distribution around zero, beyond the maximum-residual figures already reported in Tables 4–5) is left for future refinement of the real fit obtained in Section 3.1; it is not possible for the [2] dataset (Section 3.2).

Three limitations must be acknowledged. The WKB approximation for a thick barrier 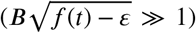 ceases to hold near *t*^*^, where a uniform description (Airy functions, analogous to the treatment of classical turning points) would be necessary to avoid the artificial abrupt transition produced by the simple formula. The uniform correction near *t*^*^ can be obtained via:

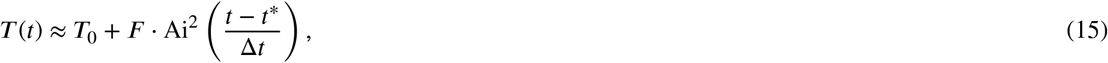

where Ai is the Airy function and Δ*t* is the width of the transition region. In addition, the adiabatic hypothesis — *f*(*t*) varying slowly on the spatial scale of the barrier — can be checked, at the order-of-magnitude level, against the fitted degradation timescales *τ*. The condition (Eq. 2) requires *τ* ≫ *π*ℏ/*E*; taking *E*at the order of the thermal energy *k*_*B*_*T* at skin/body temperature (*T* ≈ 305 K, *E* ≈ 0.026eV) as a conservative estimate for the drug molecule’s kinetic energy scale — since *E* itself is not independently constrained by the fit, only the ratios 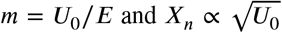 are — gives a threshold *π*ℏ/*E* ≈7.9 × 10^−14^ s. The fitted *τ* for the well-identified systems (2.688 h for G-5-FU, 4.373 h for NCA-1-5-FU, i.e.∼ 10^4^ s) exceeds this threshold by a factor of order10^17^. This margin is so large that the conclusion is insensitive to the precise, unmeasured value of *E*: even an unrealistically small *E* several orders of magnitude below thermal energy would leave the adiabatic condition comfortably satisfied. The approximation is therefore justified for the systems studied here, though this remains an order-of-magnitude argument rather than a measurement of *E* itself.

Furthermore, the quantum kinetic framework proposed by De Moura and Albuquerque (1990) opens an elegant pathway to model drug transport through macro-biological barriers, such as the cranium in transcranial drug delivery systems. Instead of matrix degradation, the time-dependent effective potential *U*_eff_(*t*) = *U*_0_ *f*(*t*) could describe the transient permeabilization of bone tissue induced by external stimuli (e.g., focused ultrasound). In this scenario, the traversal time formulation provides a strict physical metric to estimate the structural transit time of nanocarriers across dense mineralized matrices, bridging fundamental quantum mechanics and advanced neuro-pharmacokinetics.

A third limitation concerns overfitting risk: the exponential model has 5 free parameters fitted to 13–14 digitized points without replicate measurements, leaving only 8–9 degrees of freedom. The excellent fit quality reported for NCA-1-5-FU and G-5-FU (*R*^2^ > 0.99) should be read with this in mind. Leave-one-out cross-validation was considered but is of limited diagnostic value at this sample size (removing one of 13–14 points removes∼7–8% of the already sparse dataset per fold, and re-running a 5-parameter global optimization on the remainder offers little power to detect overfitting distinctly from ordinary parameter uncertainty); a more informative test would be an independent replicate dataset or additional time points for the same formulations, which we recommend as a direction for future experimental work rather than a further re-analysis of the same 13–14 points.

A connection with classic empirical models can also be established. For intermediate times, the exponential model with *f*_∞_ = 0reduces to:

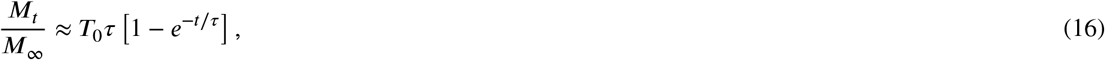

which recovers the functional form of the first-order model. On the other hand, the rational model with *n* = 1/2 approaches the Higuchi model 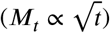 for intermediate times, showing the flexibility of the approach.

### 4.3. Connection with Partition-Controlled Models

The time-dependent barrier model developed in this work shares a direct physical foundation with the partition-controlled kinetic framework proposed by de Albuquerque and de Albuquerque [17]. In that macroscopic model, the effective rate constant is given by:

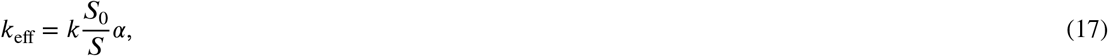

Where *S* is the equilibrium solubility in the oily core, *S*_0_ is a reference solubility, and *α* is a dimensionless correction factor [17].

Within our semiclassical framework, the observed inverse-solubility scaling (*k*_eff_ ∝*S*^−1^) finds a natural microscopic interpretation: a higher affinity and solubility *S* within the liquid core implies a stronger thermodynamic retention of the drug molecules, effectively increasing the potential barrier height *U*_0_ (and consequently the energy ratio *m* = *U*_0_/*E*). This directly reduces the instantaneous tunneling transparency *T*(*t*).

Furthermore, the correction factor *α*, which accounts for structural non-idealities in the diffusion-partition limit [17], serves as a macroscopic aggregate of our parameters *T*_0_, *F, X*_*n*_, and the matrix degradation timescale *τ*. This cross-scale consistency is strongly supported by the reanalysis of independent experimental data for adapalene nanocapsules [17]: while traditional multiplicative models fail to explain the kinetics, the inverse-solubility scaling collapses the release constants of distinct formulations (melaleuca oil vs. Miglyol®) to within a tight 6.5% variance, despite a 4.4-fold difference in absolute solubility [17]. The time-dependent potential barrier model thus provides the explicit transport dynamics that underpins these thermodynamic scaling laws.

## 5. Conclusions

We have argued that the potential barrier framework for describing drug release does not require a multifractal formalism: the equations involved are the same as the standard Schrödinger equation, and the time evolution can be introduced in a physically motivated manner via an effective barrier *U*_eff_(*t*) = *U*_0_ *f*(*t*). Rather than the thick-barrier WKB approximation used in earlier drafts, the exact transmission formula (Section 2.6) was ultimately adopted for all numerical fits, since it is smooth by construction across the barrier collapse. The two examined hypotheses for *f*(*t*)— exponential and rational (Hill) — lead to distinct and testable predictions for the barrier collapse time *t*^*^ as a function of the energy ratio *m*, providing an empirical criterion to discriminate between hypotheses.

We have established that:

1. The model with a time-dependent barrier (*U*(*x, t*) = *U*_0_*f*(*t*)) is mathematically consistent and physically motivated.
2. The exact (non-WKB) transparency was fitted, via global optimization, to real digitized *ex-vivo* permeation data [1], substantially improving fit quality over the WKB approximation for all three systems tested, and producing genuinely smooth theoretical curves (Figure 1). For two of three systems (G-5-FU and NCA-1-5-FU), the exponential-family fit was well-identified (all parameter uncertainties below 11% and 6%, respectively; *R*^2^ >0.999); the rational (Hill) family remained poorly identified for all three systems even with the improved formula, and by parsimony the exponential family is preferred throughout. For the third system (G-NCA-1-5-FU), the free-*m* fit remained non-identified given the available 13 digitized points without replicate error bars – reported as a genuine finding, not concealed. A methodological lesson worth stating explicitly: an initial, single-start fit had also reported NCA-1-5-FU as non-identified; only a global search (differential evolution) revealed that this was a local-optimum artifact, not a genuine data limitation – underscoring the importance of global, rather than local, optimization for this class of oscillatory transmission models.
3. A sensitivity check fixing *m* at the G-5-FU-derived value (*m* = 1.377) resolved the identifiability problem for G-NCA-1-5-FU at negligible cost in fit quality (Table 5, Figure 2), consistent with a shared energy ratio for that system. For NCA-1-5-FU, however, which is independently well-identified with its own, distinct value (*m* = 1.188±0.007), forcing the shared value degraded the fit by a factor of ∼ 4 in maximum residual – a small but genuine cost. The two independently-fitted values (NCA-1-5-FU: 1.188; G-5-FU: 1.377) differ by ∼ 4σ(*z* ≈ 4.0), a formally significant difference.**The data therefore do not support a single, universal energy ratio** *m***across all systems studied**, and this hypothesis is rejected for the complete set; the shared value remains consistent only with G-NCA-1-5-FU, whose own *m* cannot be independently constrained.
4. The proposed approach dispenses with fractal formalisms or artificial space-time transformations, offering a more parsimonious and generalizable theoretical framework, while acknowledging that spatially grounded fractal descriptors retain legitimate applications elsewhere in oncology [18].

On the basis of the real-data validation obtained so far (Section 3.1), the proposed framework offers a parsimonious, well-constrained description for the well-identified system (G-5-FU); direct physical interpretation of its parameters (*τ, f*_∞_, *T*_0_, *X*_*n*_), in place of phenomenological fractal constants; an experimentally testable barrier collapse time *t*^*^; and recovery of classical release models (first-order, Higuchi) as limiting cases (Section 2.3). The persistent non-identifiability of the free-*m* fit for two of the three systems tested, and its partial resolution via the shared-*m* sensitivity check, are themselves informative results about the data requirements and inter-formulation structure of this class of model.

Future work should: (i) obtain additional digitized time points (concentrated in the 0–5 h window) or, preferably, replicate measurements with per-point error bars for NCA-1-5-FU and G-NCA-1-5-FU, to independently confirm the shared-*m* hypothesis rather than relying on the sensitivity check of Section 3.1; (ii) replace the order-of-magnitude thermal-energy estimate of *E* used in the adiabatic check above with an independent value (e.g. from molecular dynamics or an explicit choice of barrier width *a* and effective mass *μ*), tightening the validation of the adiabatic hypothesis underlying the traversal-time expressions of Section 2.4; (iii) apply the uniform Airy-function correction near *t*^*^ for any residual sharp features at extreme parameter values; (iv) seek unpublished raw time-course data from the authors of [2] for their four systems, since only a single 24 h endpoint per system is available in the published text (Table 1), precluding an independent multi-parameter fit; (v) test the predictions against the additional aptamer-functionalized systems identified in the Introduction [5–7]; and (vi) extend the model to arbitrary barrier profiles (triangular, parabolic), stochastic/thermal effects in *f*(*t*), and a hybrid tunneling-diffusion formalism for long-term release regimes.

## Acknowledgments

None

## CRediT authorship contribution statement

**Maria Antonia S. de Albuquerque:**Conceptualization, Methodology, Investigation, Writing – original draft, Writing – review & editing.

**Douglas F. de Albuquerque:**Conceptualization, Methodology, Formal analysis, Investigation, Writing – original draft, Writing – review & editing, Visualization.

## Data availability

No new experimental data were generated. The 24 h release efficiencies of Table 1 are taken directly from the published studies [2–4]. The *ex-vivo* permeation curves of Section 3.1 were digitized from Figure 3 of [1] using Engauge Digitizer and cross-checked against the maximum permeated amounts quoted in that paper’s text (agreement within 1.5%).

The digitized experimental coordinates used for the non-linear regression, along with the complete Python implementation of the fitting procedure described in Section 3.1 (including the exact transmission formula of Section 2.6, the global-optimization routine, and the residual analysis), are available as Supplementary Material upon request to the corresponding author.

## Institutional review board statement

Not applicable.

## Informed consent statement

Not applicable.

## Abbreviations

5-FU: 5-Fluorouracil
AS1411: Anti-nucleolin DNA aptamer used for tumor targeting
NCA-1: Aptamer-functionalized polymeric nanocapsule sample [4]
L4: Aptamer-functionalized liposome sample (L4Apt-5FU-15) [3]
C1: Oil-in-water cream base formulation [2]
WKB: Wentzel–Kramers–Brillouin (semiclassical) approximation
*U*_eff_(*t*): Time-dependent effective barrier height
*t*^*^: Barrier collapse time
*m*: Energy ratio,*U*_0_/*E*

